# Full resolution HLA and KIR genes annotation for human genome assemblies

**DOI:** 10.1101/2024.01.20.576452

**Authors:** Ying Zhou, Li Song, Heng Li

**Affiliations:** Department of Data Science, Dana-Farber Cancer Institute, Boston, MA, 02115, USA; Department of Biomedical Data Science, Dartmouth College, Hanover, NH, 03755, USA; Department of Biomedical Informatics, Harvard Medical School, Boston, MA, 02115, USA

**Keywords:** HLA haplotype, KIR haplotype, long-reads assembly

## Abstract

The HLA (Human Leukocyte Antigen) genes and the KIR (Killer cell Immunoglobulin-like Receptor) genes are critical to immune responses and are associated with many immune-related diseases. Located in highly polymorphic regions, they are hard to be studied with traditional short-read alignment-based methods. Although modern long-read assemblers can often assemble these genes, using existing tools to annotate HLA and KIR genes in these assemblies remains a non-trivial task. Here, we describe Immuannot, a new computation tool to annotate the gene structures of HLA and KIR genes and to type the allele of each gene. Applying Immuannot to 56 regional and 212 whole-genome assemblies from previous studies, we annotated 9,931 HLA and KIR genes and found that almost half of these genes, 4,068, had novel sequences compared to the current Immuno Polymorphism Database (IPD). These novel gene sequences were represented by 2,664 distinct alleles, some of which contained non-synonymous variations resulting in 92 novel protein sequences. We demonstrated the complex haplotype structures at the two loci and reported the linkage between HLA/KIR haplotypes and gene alleles. We anticipate that Immuannot will speed up the discovery of new HLA/KIR alleles and enable the association of HLA/KIR haplotype structures with clinical outcomes in the future.

## Introduction

The HLA genes and KIR genes play a central role in immune responses. They are clustered in two genomic regions, respectively. The HLA genes are located in the Major Histocompatibility Complex (MHC) region on the short arm of chromosome 6. The MHC region spans about 5 megabase (Mb). Coding genes in MHC, such as *HLA-A*, *B*, *C*, *DR*, *DP*, and *DQ* are highly polymorphic among human populations, which is maintained by balancing selection (DeGiorgio et al. 2014; Meyer et al. 2018). Many HLA Alleles (Marsh et al. 2010), or sub-groups distinguished by gene sequences, were reported to be associated with hundreds of immune-related diseases (Gough and Simmonds 2007; Medhasi and Chantratita 2022), and HLA allele matching (*HLA-A*,*B*,*C*,*DRB1*, and *DQB1*) has been widely used in pairing unrelated donor and recipient to lower the failure risk of graft transplantation (Buck et al. 2016; Tiercy 2016; Dehn et al. 2019).

The KIR gene cluster is a smaller region located in chromosome 19, ranging from 150 kilobase (kb) to 200kb in different individuals. It contains 17 genes, including 15 coding genes and 2 pseudo-genes, regulating the development, tolerance and activation of natural killer (NK) cells. Different from HLA genes’ high polymorphism, KIR genes share similar gene structures and with diverse copy numbers for each haploid genome (Carrington and Norman 2023). The most common KIR haplotype is called the “A-haplotype” with 9 genes, and all other KIR haplotypes are defined as the “B-haplotype”(Cisneros et al. 2020). The A-haplotype is also called the inhibitory haplotype because most of its KIR genes inhibit the acBvaBng of NK cells, while the B-haplotype tends to cover more activating genes. Similar to the HLA genes, KIR genes also have a nomenclature system to functionally distinguish each KIR allele sequence (Robinson et al. 2018).

High-throughput sequencing with short reads has become a common approach to typing HLA/KIR alleles, which enables discovering the connection between special gene groups and clinical outcomes in large cohort(Sakaue et al. 2023). By searching the best combinations of alleles that explained by the reads data, many computational methods can achieve high sensitivity and accuracy when the gene has been well documented (Szolek et al. 2014; Norman et al. 2016; Kawaguchi et al. 2017; Lee and Kingsford 2018; Dilthey et al. 2019; Sverchkova et al. 2019; Kim et al. 2019; Gao et al. 2022; Song et al. 2023). For example, the classic HLA genes (*A*, *B*, *C*, *DRB1*, *DQB1*, *DPB1*) have hundreds to thousands of allele sequences in the IPD-IMGT/HLA database (Barker et al. 2023). However, for underrepresented non-classical HLA genes, these tools would have reduced power, and some of them would not even report those genes.

Genotyping with long reads may overcome this limitation as each gene sequence can be fully assembled, which will lower the requirement of reference gene sequences. Thus, it is more straighjorward to annotate and call alleles on an assembly rather than from raw reads data. Recently, more and more long reads full genome assemblies have been released. For example, the Human Pangenome Reference Consortium (HPRC) released 94 haploid assemblies in the phase I (Liao et al. 2023), the number will go up to several hundred in the phase III by mid-2024. The Chinese Pangenome Consortium (CPC) released 114 haploid assemblies in the phase I (Gao et al. 2023). There is an urgent need to study HLA/KIR regions on those assemblies.

To this end, we developed a new computation tool, Immuannot, to annotate HLA and KIR gene structures on human assemblies and output the allele calls at the same time. Taking IPD-IMGT/HLA and IPD-KIR as references, Immuannot employs full gene sequences to resolve gene structures and uses all available coding sequences (CDSs), including the ones with partial CDS, to type alleles. We applied our pipeline to 212 full genome assemblies, 50 KIR reference contigs, and 6 HLA reference genome. Immuannot reported thousands of novel HLA and KIR allele sequences that were missing in the IPD database, particularly for those non-classical coding genes and pseudo-genes. Through a series of genetic analyses, we validated Immunannot and identified numerous new genetic patterns. As long-read data become more and more popular, our tool will improve the genetic research in HLA and KIR regions.

## Result

### Method overview

Immuannot types 36 HLA genes in the IPD-IMGT/HLA data v3.52.0 and 17 KIR genes in the IPD-KIR database (V2.12) (Barker et al. 2023; Robinson et al. 2013). It first aligns gene sequences, including introns, to contigs of interest (Figure 1A). A gene is initially detected on a contig if 90% of the gene sequence is mapped. When multiple reference allele sequences match the same location of the target contig, the one with the largest gap-compressed identity and longest matching length is chosen as the template allele, which is further used for determining gene structure, such as exon boundaries, and allele type. If the template allele has a perfect match to the contig, the contig is reported to carry that allele. Otherwise, CDS is extracted for allele calling with extra steps. We use alleles with full gene length sequences for template selection and use all alleles with full or partial CDS for allele typing.

**Figure 1:**
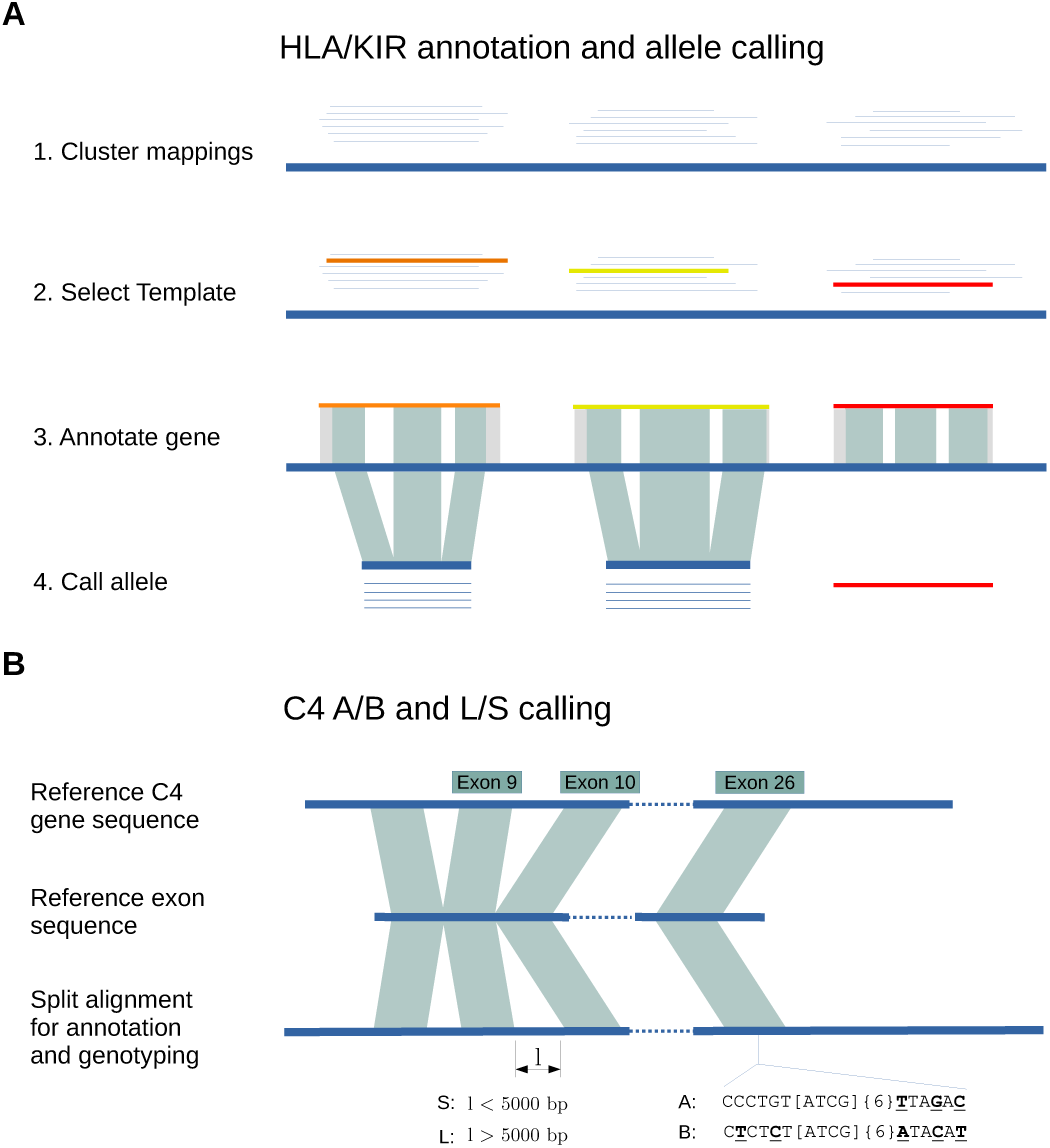
Overview of the Immuannot algorithm. (**A**) Annotate HLA/KIR genes using known alleles from the IPD-IMGT/HLA or the IPD-KIR databases. Gene sequences (thin line) from the IPD databases are mapped to the target contig sequence (thick line) at step 1. At step 2, the best-matching allele with the smallest number of mismatches and the longest alignment length is chosen as the template allele (in different colors) for each mapping cluster. Project the gene structure of template allele to the contig to annotate exons, introns, UTRs, CDS, and start and stop codons (step 3). At step 4, determine the allele type based on the CDS similarity. (**B**) Annotate the C4 genes. Align the C4 transcript sequence from GRCh38 to the contig sequence. Infer C4A vs C4B based on the sequence on exon 26 and infer the long- vs short-form based on the intron size between exon 9 and 10.

In order to annotate complement component 4 (*C4*), a MHC class III gene that is not covered by the IPD-IMGT/HLA database, we apply a different strategy (Figure 1B). In this case, exons are extracted from one reference gene sequence and aligned to the target contig with splicing. We can annotate the gene structure, and identify the length of the 9th intron and the key peptides in the 26th exon to infer the long- vs short-form and C4A vs C4B types (Yu et al. 1986; Blanchong et al. 2001; Wang and Liu 2021).

Immuannot can distinguish multiple copies of a particular gene because each reference allele sequence is allowed to be mapped to multiple regions/contigs. It outputs in the GTF format, including the gene structure annotations and the allele type defined by the nomenclature system (Marsh et al. 2010; Robinson et al. 2018). When a novel allele is detected, we will provide a consensus call with a ‘new’ tag in the corresponding naming field to summarize the effects of new nucleotide variants. More details about our pipeline can be found in the Methods section.

Notably, IPD-IMGT/HLA and IPD-KIR include both protein-coding genes and pseudogenes. In their master annotation file, they provide “CDS” coordinates for pseudogenes but without translated proteins due to frameshifts or in-frame stop codons in the CDS. We could not find out how CDSs for pseudogenes were determined. Either way, following the IPD-IMGT/HLA and IPD-KIR annotations, Immuannot reports CDS for all genes as well, including both protein-coding genes and pseudogenes.

### Evaluation

To evaluate our pipeline, we first applied Immuannot to eight well-annotated MHC assemblies that were published in 2023 (Houwaart et al 2023). We compared our allele calls to theirs on HLA genes and C4 genes. Among 72 full-resolution comparisons of HLA protein-coding genes (*A*, *B*, *C*, *DRB3*/*4*/*5*, *DRB1*, *DQA1*, *DQB1*, *DPA1*, *DPB1*) across eight haplotypes (Supplemental Table S1), we only observed one inconsistency with the calls by Houwaart et al. In the sample OK649235, we called *03:01:01:02* for the HLA-DRB1 gene, while Houwaart et al called *03:01:01:01*, different in the fourth field. Further investigation with alleles from IMGT database shows that allele *DRB1*03:01:01:02* is perfectly aligned to the assembly OK649235 while allele *DRB1*03:01:01:01* has 11 mismatches (a 10bp deletion and a substitution) in introns. Our C4 typing is identical to the results by Houwaart et al as well (Supplemental Table S1).

We also compared Immuannot with the recently released software SKIRT (Hung et al. 2023) for KIR allele annotation on assembled genomes. We tested both software on HPRC assemblies. On average, Immunannot took about 4 minutes to annotate KIR alleles on one full genome haploid assembly with three threads, while SKIRT spent about 24 minutes with three threads. SKIRT was slower because it heavily relied on minimap2 v2.17 (Li 2018) with split-alignment mode which was more Bme-consuming than the non-split mode. Immuannot annotated 901 genes that were all annotated by SKIRT and 99.7% (898/901) of them shared the same location and allele name. For the three mismatches, SKIRT called them as gene fusions, while Immuannot only annotated these genes without specific allele groups. SKIRT annotated five more genes not reported by Immuannot. Three of them have partial deletions with the remaining regions covering <90% of the reference sequence, and the other two include large intronic indels that fragmented the alignment and failed our template selection step (Supplemental Notes).

We further applied Immuannot and SKIRT to 50 KIR contigs which were part of the GRC38.p14 release for a three-way comparison (Supplemental Table S1). Immuannot annotated 427 genes, where 425 of them perfectly matched the SKIRT annotation. Of the two differences, the first at NW_003571057.2:827290-839484 was annotated by SKIRT as a fusion of genes *KIR2DL3* and *KIR2DP1*, but Immuannot reported as *KIR2DP1* without allele group information. As to the second difference, Immuannot found gene *KIR2DL1* at NW_016107303.1:71161-85920, which was missed by SKIRT but was confirmed by the GRC annotation. SKIRT identified 18 additional genes that were not found by Immuannot. We manually checked six cases and found all of them to be partial genes. Among these 18 genes, 12 were reported by GRC and 11 of them had identical annotations. GRC annotated two genes not reported by either SKIRT or Immuannot. Both genes were annotated as *KIR3DX1*, which was absent from the IPD-KIR database (Supplemental Table S1).

### Novel alleles

We applied Immuannot to 212 full genome haploid assemblies, including HPRC (n=94) (Liao et al. 2023), CPC (n=114) (Gao et al. 2023), CHM13.0 (Nurk et al. 2022), GRCh38.p14 (Schneider et al. 2017), and CN-T2T (n=2) (He et al. 2023), 50 KIR reference contigs and 6 HLA reference assemblies (Houwaart et al. 2023). This resulted in 7,519 HLA genes, represented by 2,673 unique alleles as some unique alleles were present in multiple samples. Furthermore, 2,696 of these genes (1,531 unique alleles) were not seen in the IPD-IMGT/HLA database. As to KIR, Immuannot annotated 2,412 genes, represented by 1503 unique alleles, and 1,372 of these genes (1,133 unique alleles) were novel. In total, 41.0% [(2696+1372)/(7519+2412)] of HLA/KIR genes had novel alleles absent from the IPD databases.

In more detail, among the six classical HLA protein-coding genes (*HLA-A*, -*B*, -*C*, -*DRB1*, -*DPB1*, and -*DQB1*), 22.1% (289/1310) genes annotated by Immuannot were novel but only 2.4% (7/289) carried mutations that led to amino acid changes (Supplemental Table S2). All seven novel proteins had distinct sequences. Among other HLA protein-coding genes (i.e., not pseudogenes), 31.3% (1080/3454) were novel and 10.2% (110/1080) had amino acid changes. These 110 protein-coding genes were represented by 43 unique alleles. At the KIR locus, 57.8% (1116/1931) KIR protein-coding genes were novel, and 4.2% (47/1116) had amino acid changes, represented by 42 novel unique protein sequences (Figure 2, Supplemental Table S2). Among pseudogenes in the IPD databases, 48.9% (1583/3236) were novel.

**Figure 2:**
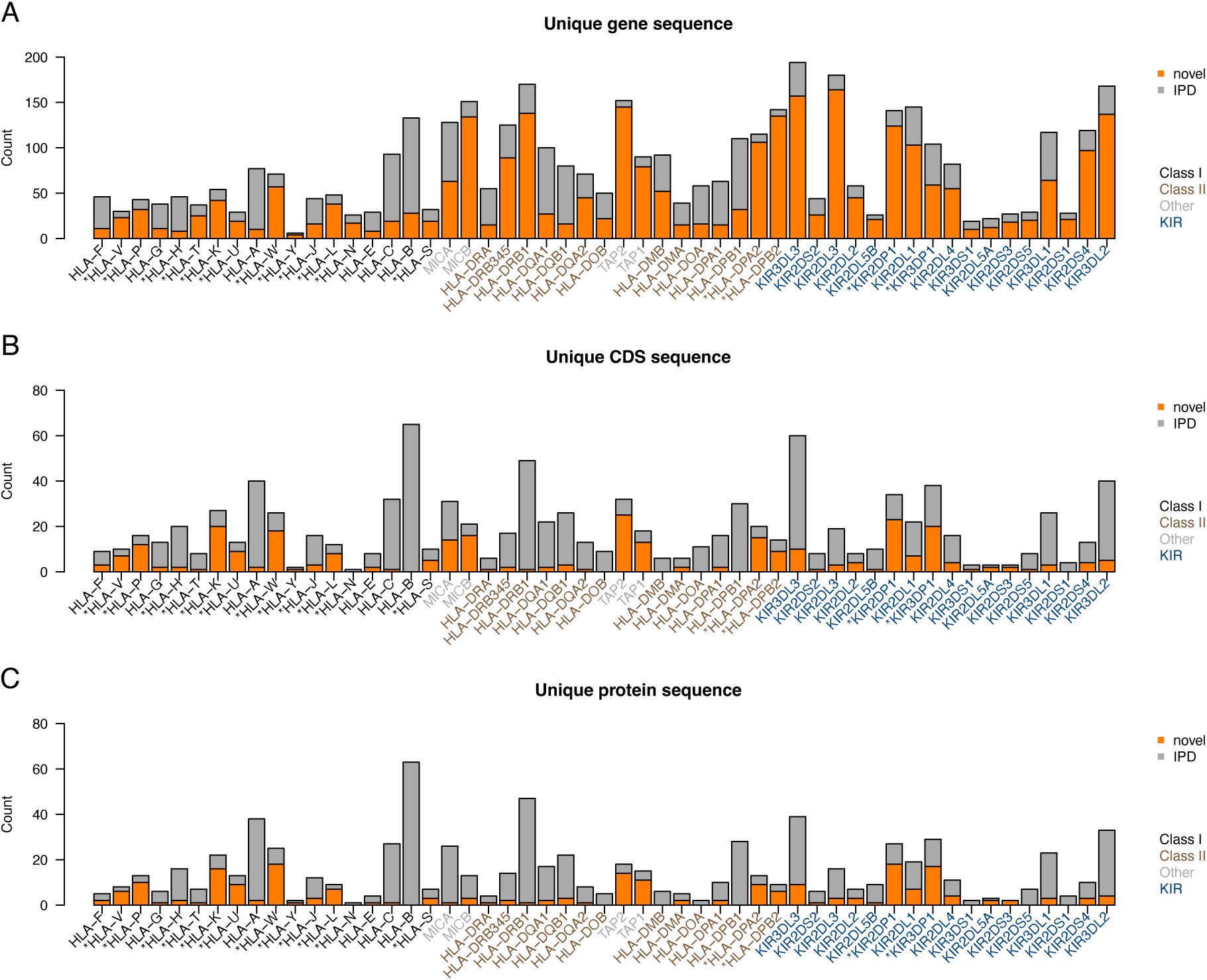
Summary of unique allele sequences. Alleles with identical sequences are merged. Known allele sequences in the IPD databases are shown in gray and novel allele sequences are shown in orange. (**A**) Gene sequences with intron included. (**B**) CDS sequences. (**C**) Protein sequences.

### HLA Gene diversity

We investigated the CDS diversity of HLA and KIR genes (Figure 3). *HLA-DQB1* had the highest diversity in our dataset with the average pairwise sequence divergence approaching 5% (Figure 3B). Several other well-studied HLA genes, such as *HLA-A*, *-B*, *-C* and *-DRB1* were also highly diverse, whose average pairwise sequence divergence is over 2.5% among either Africans in HPRC or Asians in CPC.

**Figure 3:**
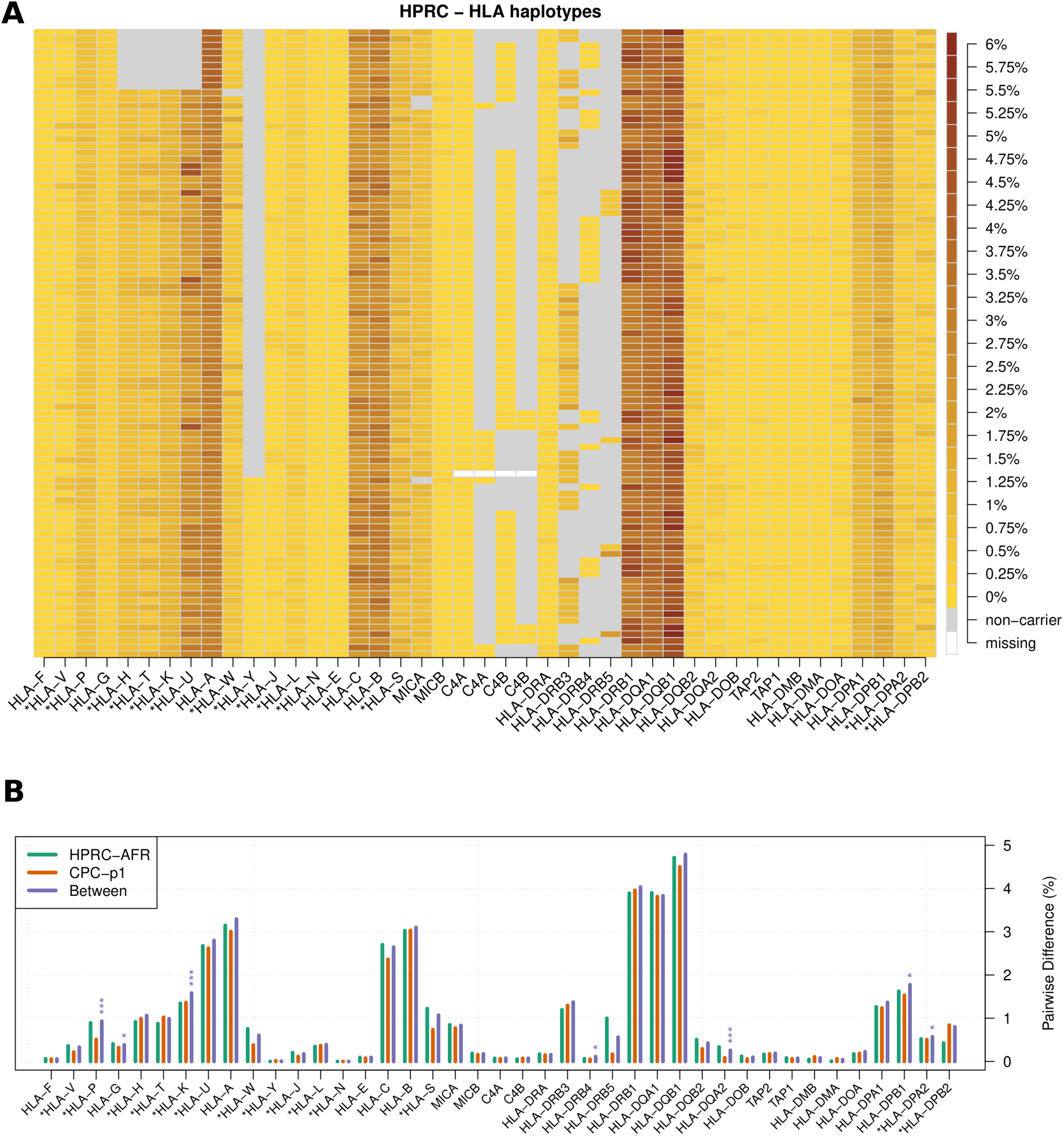
The diversity of HLA coding sequences. (**A**) HLA haplotypes. Each row represents one HLA haplotype. For each haplotype, heat from light yellow to the dark brown indicates the average pairwise CDS divergence between the haplotype and all the other haplotypes. A gray box denotes the deletion of a gene in the middle of a contig, while a white box denotes an unassembled gene in assembly gaps. (**B**) HLA within-population diversity and between-population divergence. Each group of bars shows average pairwise CDS difference within HPRC African samples (AFR, n_hap = 24), within CPC Asian samples (n_hap = 114), and between the two populations. Permutation tests were performed to test whether the two populations are distinguishable in terms of CDS divergence. P-values are adjusted with Bonferroni method. Number of stars indicates different significance level: ’*’ for p-value < 0.05, ’**’ for p-value < 0.01, ’***’ for p-value < 0.001.

There are several class-I pseudogenes. Among them, *HLA-U* had the highest diversity with pairwise sequence divergence similar to *HLA-A*’s. Despite being a pseudogene, *HLA-U* is also expressed in tissues such as whole blood, lung, and EBV-transformed lymphocytes (Supplemental Figure S2). On the other hand, *HLA-Y* and *HLA-N* had the lowest diversity. *HLA-Y* is a pseudogene possibly derived from *HLA-A* (Williams et al. 1999). It was in 28.7% of HPRC haplotypes and 12.7% of CPC haplotypes, and the CDS diversity was 0.0% in HPRC and 0.017% in CPC among all carriers. *HLA-N* is first reported by Geraghty et al (Geraghty et al. 1992a). It is annotated with one exon and the CDS region has only 161bp. All HPRC and CPC samples were found to carry identical CDS, and the estimated CDS diversity is 0.002% among samples from the 1000 Genomes Project (n=3202; Supplemental Table S3).

In most genomic regions outside HLA, African populations have higher diversity than non-African populations due to the out-of-Africa bottleneck in non-Africans populations(1000 Genomes Project Consortium et al. 2015; Mallick et al. 2016; Bergström et al. 2020). Nevertheless, African samples in HPRC (n=40) and Chinese samples in CPC (n=57) have similar diversity for most HLA genes (Figure 3B). The pairwise CDS sequence divergence between the two populations is also indistinguishable from within-population pairwise divergence based on a permutation test (Methods). This observation is probably caused by balancing selection that tends to maintain the high diversity in MHC.

### HLA gene deletions

Deletion of *HLA-Y* (*Y*-del) is the most common gene deletion among HLA genes (71.3% in HPRC and 87.3% in CPC). It is rarely studied in large cohorts because the human reference genome, such as the release of GRCh38, does not contain this gene. With long-read assembled haplotypes, we found that *HLA-Y* was highly associated with specific HLA-A allele groups:

*A*29,30,31,33* and *68* (Figure 4A). This association was especially strong in Han Chinese samples. (Supplemental Figure S3B)

**Figure 4:**
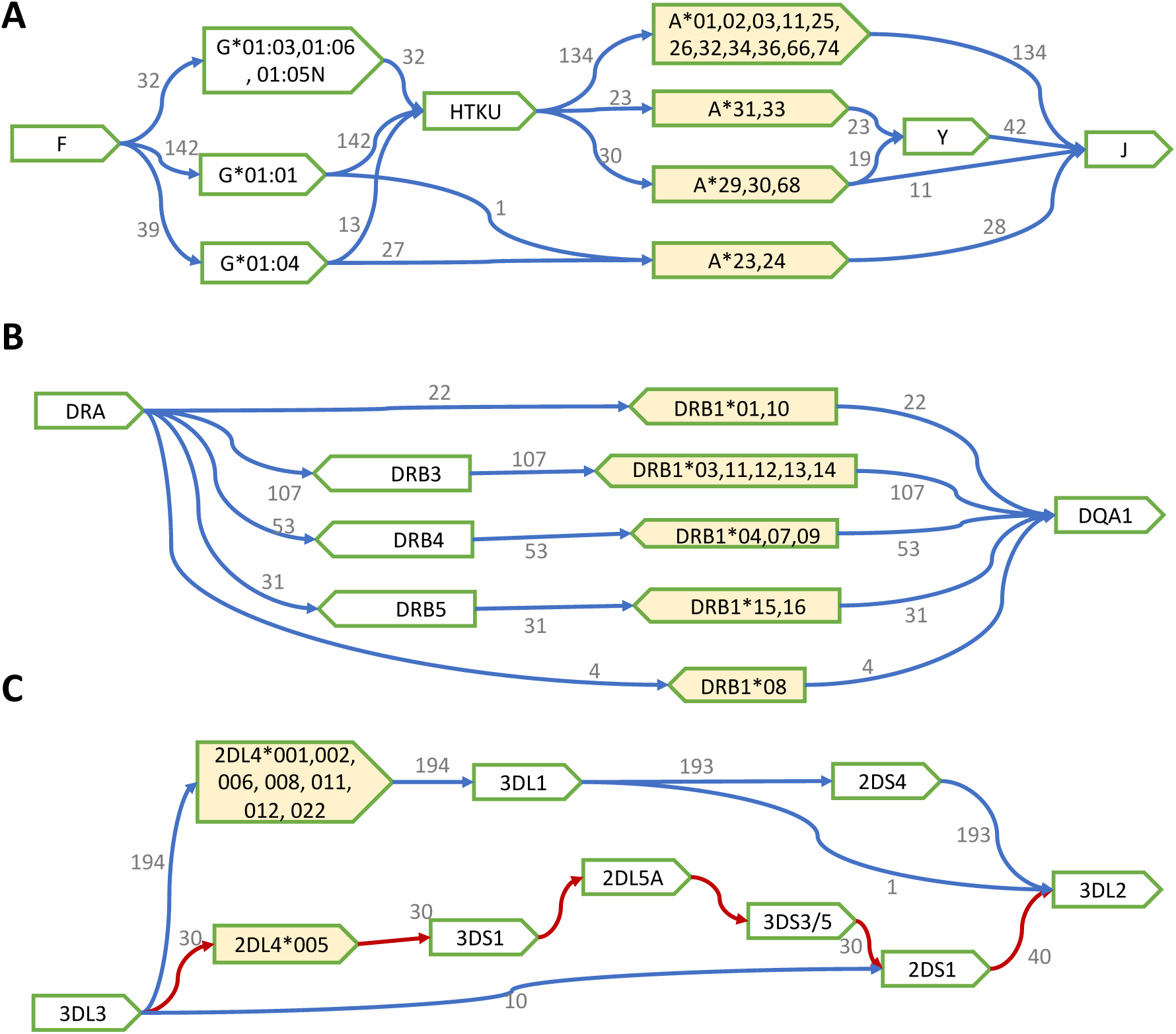
Local haplotype structures and their linkage to HLA/KIR alleles. (**A**) Haplotypes with the deletion of the *HLA-H*, *T*, *K* and *U* genes (*HTKU-del*) and the deletion of the *HLA-Y* gene. (**B**) DR haplotypes. (**C**) Haplotypes in the KIR telomere region.

The second most common HLA gene deletion we observed was the deletion of four pseudo-genes: *HLA-H*, *T*, *K* and *U* (*HTKU-del*), which was reported to be in linkage disequilibrium with *HLA-A*23/24* and *HLA-G*01:04* (Geraghty et al. 1992a, 1992b; Paganini et al. 2019; Carlini et al. 2016). In our data, *HTKU-del* happened in 9.6% HPRC assemblies and 16.4% in CPC assemblies, and it was highly linked to *A*23/24*. Different from earlier observations, we also found 13 *G*01:04* haplotypes had HTKU genes, and 27 *G*01:04* haplotypes had *HTKU-del* (Figure 4A). We observed that all *HTKU-del* haplotypes had *Y-del*, but some *Y-del* haplotypes still carried the four pseudo-genes, suggesting that *HTKU-del* event might be more recent than *Y-del*. Using the read depth information, we genotyped *HTKU-del* events for samples from the 1000 Genomes Project (Byrska-Bishop et al. 2022) (Supplemental Figure S4). The estimated allele frequency of *HTKU-del* was 39.4% in the Japanese population (JPT), 18.8% in Chinese (CHS + CHB), and 14.3% in African and American together.

Protein-coding gene *MICA* may also be deleted. *MICA* functions as a stress-induced ligand for integral membrane protein receptor *NKG2D* (Bauer et al. 1999). The deletion of *MICA* has low frequency in HPRC (3.2%) and CPC (1.9%) but has higher frequency in a Mexican population (MXL, 11.9%) and in a Peru population (PEL, 5.7%) (Supplemental Figure S5). Additionally, we also found that around 22.1% of all our *MICA* calls have incomplete ORF (Supplemental Table S3).

### HLA DR haplotypes

The HLA class II region is highly structured (Klitz et al. 2003; Norman et al. 2017; Chin et al. 2023), which is commonly characterized by DR haplotype. Following earlier work (Houwaart et al. 2023), HLA DR haplotype is defined by the combination of *HLA-DRB1* allele groups and the presence or absence of DRB3/4/5 genes. All 217 HLA haplotypes in our analysis are in agreement with DR haplotype definition: *DRB1*03*, *11*, *12*, *13*, and *14* are always linked to DRB3 (DR3), *DRB1*04*, *07* and *09* linked to DRB4 (DR4), *DRB1*15* and *16* are linked to DRB5 (DR5), and *DRB1*01*, *10*, or *08* never carry DRB3/4/5 alleles (DR1+DR8) (Figure 4B).

We further explored the polygenetic relationship of DRB1/3/4/5 alleles based on CDS differences (see Methods). The DRB1 alleles from each of the DR3, DR4, and DR5 groups formed a separate clade, but the DRB1 alleles from the DR1 group were split into two different clades, suggesting two different evolutionary paths leading to the formation of the DR1 haplotypes in human (Supplemental Figure S6).

### C4 genotypes

Immuannot also annotates one MHC class III gene, complement component 4 (C4). A C4 gene consists of 41 exons, encoding a transcript of 5.4 kb in length. The human reference genome GRCh38 has two C4 genes, *C4A* and *C4B*, which have different chemical reactivities and binding affinities in interacting with antigens (Yu et al. 1986; Blanchong et al. 2001; Wang and Liu 2021). Both *C4A* and *C4B* may have two forms: long form (L) of 20.6 kb in length on GRCh38, or short form (S) of 14.6 kb. The long-form C4 gene harbors a 6.2k insertion of endogenous retrovirus sequences in the 9^th^ intron while the short-form C4 gene does not (Yu et al. 1986; Blanchong et al. 2001). Therefore, there are four possible types of C4 genes: *C4AL*, *C4AS*, *C4BL* and *C4BS*. Their copy numbers vary in different individuals and the copy-number changes may result in different disease risks (Dodds et al. 1996; Blanchong et al. 2001; Jüptner et al. 2018; Yang et al. 2004; Jaatinen et al. 1999; Breukink et al. 2015; Sekar et al. 2016). Immuannot annotates all four types of C4 genes.

In our data, *C4A* achieved a frequency of 57%. Every haplotype had at least one copy of *C4A* gene, and no haplotypes had more than two *C4A* genes or more than two *C4B* genes. The frequencies of monomodular (one copy of C4), bimodular (two copies of C4), and trimodular (three copies) were 13.9%, 79.8%, and 6.3%, respectively, for HPRC, and 8.8%, 71.9%, and 19.3% for CPC. No quadrimodular case (four copies of C4) was detected (Supplemental Table S4).

Human *C4A* and *C4B* genes are more than 99% identical. In the phylogenetic tree constructed from the CDS of all C4 genes sequences in our data, *C4A* and *C4B* were clearly separated, but the long and short forms were mixed in both *C4A* and *C4B* clades (Supplemental Figure S7). In addition to the five SNPs in exon 26 that are often used to distinguish *C4A* and *C4B*, four SNPs in exon 28 can also separate most, though not all, *C4A* and *C4B* genes. The imperfect linkage suggests possible gene conversions or recombination between exon 26 and exon 28 (Supplemental Table S4).

### KIR haplotypes

We collected 234 full-length KIR haplotypes that defined by including both *3DL3* and *3DL2* in one contig. We observed 25 unique KIR haplotypes, including the most common A-haplotype at 66.2% frequency and 24 B-haplotypes. The B-haplotypes may contain 5-15 KIR genes, including 1-5 activator genes in addition to *2DL4*. We also observed the deletion (n=10) and duplication (n=6) of two framework genes, *3DP1* and *2DL4* (Figure 5). Among the 44 samples with both KIR haplotypes fully assembled in HPRC, 18 carry two A-haplotypes (AA carriers), 21 were AB carriers, and only 5 were BB carriers.

**Figure 5:**
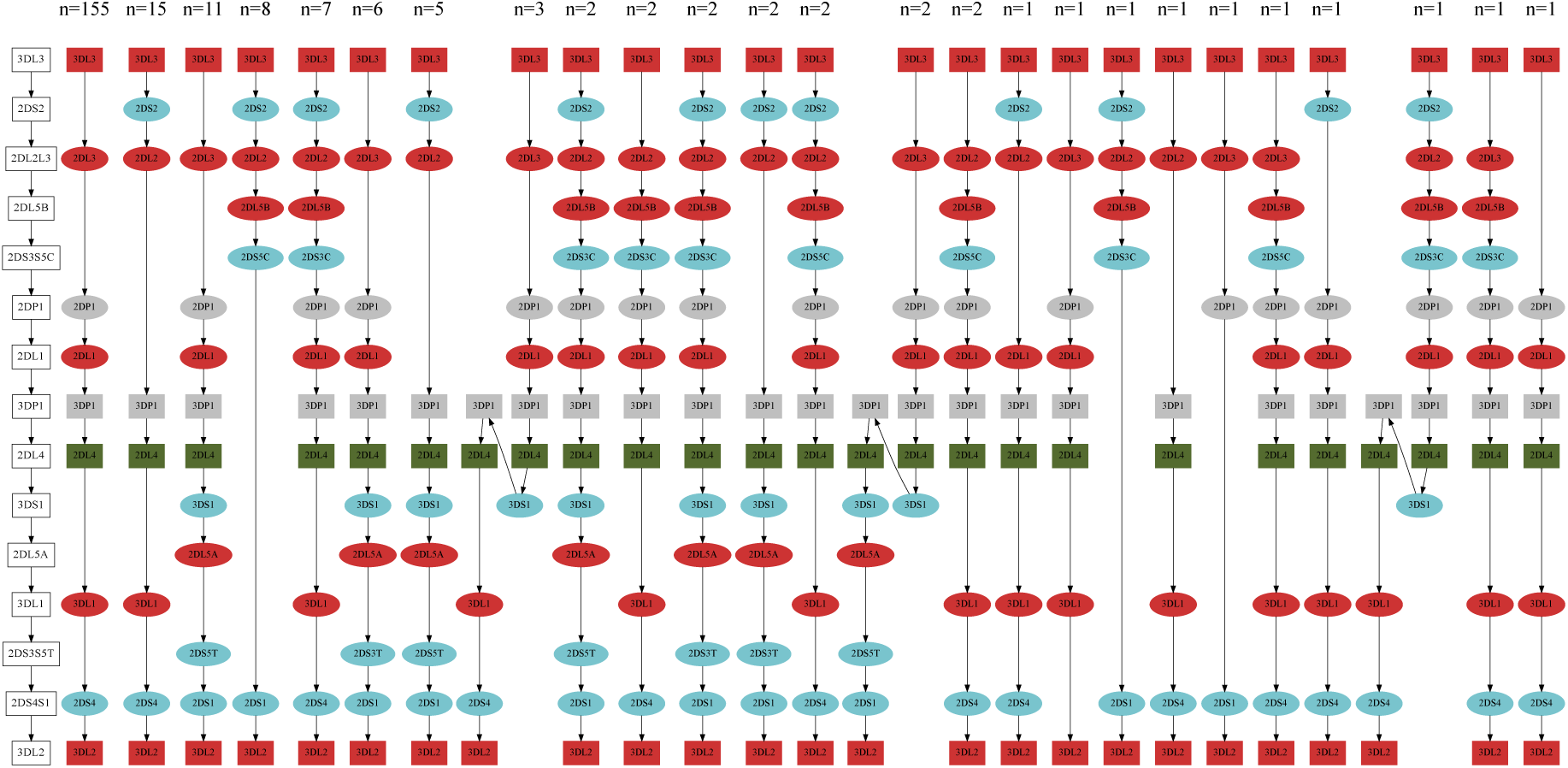
KIR haplotypes. Shape: box for framework genes and ellipse for non-framework genes. Color: red for inhibitor genes, blue for acLvator genes, gray for pseudogenes, and green for genes that can be both inhibitor and activator.

Due to frequent recombination between gene *3DP1* and *2DL4* (Cisneros et al. 2020), we further split the KIR region into centromere region (c-region, from *3DL3* to the first *3DP1*) and telomere region (t-region, from the last *2DL4* to *3DL2*). We observed 15 unique haplotypes in the c-region and 11 of them were rare (≤2 copies); we observed five unique haplotypes in the t-region and one of them was rare (≤2 copies). The common c-region haplotypes and common t-region haplotypes (>2 copies) explained more than 95% of all KIR haplotypes as we observed (Supplemental Figure S8). Interestingly, the gene *2DL4* in B-haplotypes always had the allele *2DL4*005* (Figure 4C), in line with previous studies (Amorim et al, 2021).

### Evaluating T1K genotyping with short reads

There are 40 HPRC samples that also have short-read WGS data from the 1000 Genomes Project. We took the Immuannot annotation as the ground truth to evaluate the performance of genotypers that use short reads only. We in particular focused on the results from T1K (Song et al. 2023). We did not evaluate other genotypers because most of them only work for classical HLA genes, could not genotype KIR genes, and have been evaluated in other benchmark studies (Klasberg et al. 2019; Liu et al. 2021; Thuesen et al. 2022; Yu et al. 2023).

We carefully evaluated T1K for each of HLA and KIR genes (Figure 6), and also summarized in different categories (Supplemental Table S5). For classical HLA genes (*HLA-A*, *B*, *C*, *DRB1*, *DQB1*, *DPB1*) that were well curated in IPD-IMGT/HLA, T1K achieved an average sensitivity of 97.3% and precision of 98.1–98.5%, where the precision had a range of values due to different evaluation strategies for novel alleles (see Figure 6 caption). For all HLA genes, the average sensitivity of T1K is 95.5% and the precision is 87.0–97.5%. The decreasing precision was probably related to novel alleles not in the IPD database. For KIR genes, T1K achieved a sensitivity of 91.2% and precision of 83.4–89.0%, lower than the accuracy for HLA genes. This could be caused by the high copy-number variations in KIRs that were not directly modeled in T1K’s genotyping algorithm. When ignoring the copy numbers, i.e. just match the allele prediction, the sensitivity and precision increased to 93.8% and 88.0-93.2%, respectively. In sum, our benchmark suggested that genotyping with short reads yielded highly accurate results for classical HLA genes, but remained a challenge for other highly polymorphic genes.

**Figure 6:**
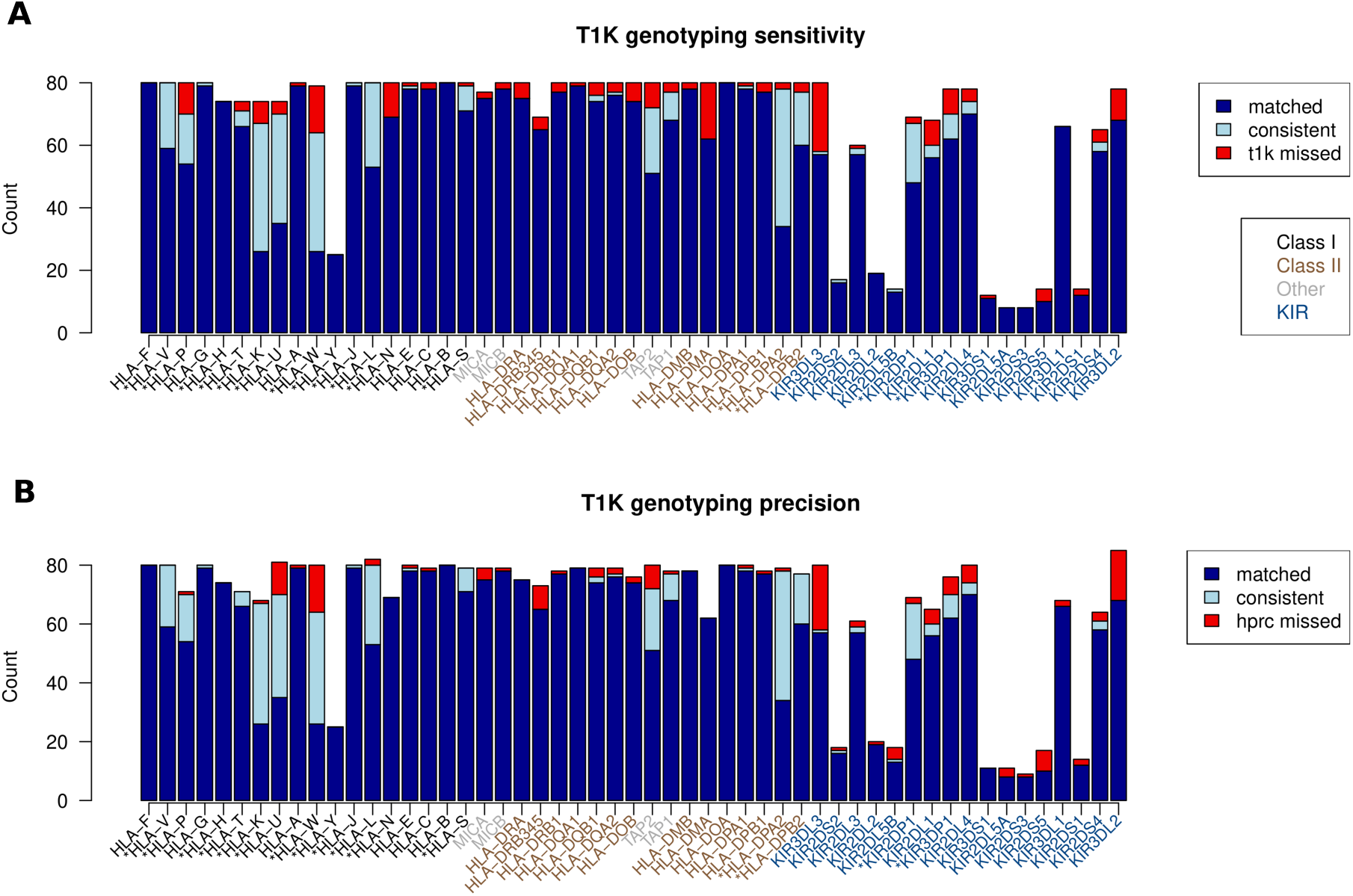
T1K evaluation for HLA/KIR genes. (**A**) Sensitivity of T1K on HPRC samples. Immuannot HPRC annotation is taken as the ground truth. An Immuannot allele is “matched” by T1K if both tools identify the allele indicating the same protein sequence. Synonymous or non-coding differences are ignored. An Immuannot allele is “consistent” with T1K if Immuannot finds a novel allele that the protein sequence not present in the IPD databases and T1K calls the correct allele group or the correct gene if no proper allele group is found in the IPD database. Different from the original evaluation in T1K paper, copy number of homozygous calls are also considered in this comparison (See Methods section for estimating copy number from T1K’s call). Otherwise, the Immuannot is “missed” and counted as an error in T1K. If Immuannot annotates a gene to have two copies but T1K only finds one, this is also considered a “missed” call. (**B**) Precision of T1K.

## Discussion

Immuannot is a convenient and efficient computation tool to annotate both HLA and KIR genes on assembled contigs at full resolution. It provides the coordinates and structures of these genes, types known gene alleles in the IPD-IMGT/HLA and the IPD-KIR databases, and also reports novel alleles. The Immuannot annotations are highly consistent with existing HLA and KIR annotations in previous work.

At present, Immuannot does not report incomplete genes caused by assembly gaps. This issue will be alleviated with better assembly. Immuannot may also miss a gene if a long insertion/deletion on a novel allele fragments minimap2 alignment. This issue can be addressed with more complete IPD databases. Jointly considering CDS alignment at the template finding step may help to alleviate the problem as well. In addition, Immuannot can only annotate HLA and KIR genes in current release, but it provides an efficient framework to conduct assembly-based annotation/genotyping when multiple reference gene sequences are presented. For example, PharmGKB (Hewett et al. 2002) collected 13 clinically important genes covering 1906 unique sequences representing different alleles. It would be interesting to know which alleles are carried by the target genome assembly for pharmacogenetics studies.

Even though HLA and KIR regions have been studied for decades, many HLA or KIR genes may not be well understood due to challenges in allele typing, and they may benefit from further study in large cohorts. For example, many south Americans lack *MICA* genes but few Europeans have this deletion. It is important to further check whether it is an adaptation process or random drift in South American. Thanks to new long-read data types and more sophisticated assembly algorithms, high-quality phased assembly will become a norm. We believe that Immuannot will provide high-fidelity HLA and KIR annotations to improve the completeness of IPD-IMGT/HLA and IPD-KIR, and to enhance the research on these genes in general.

## Methods

### Identifying candidate genes

Immuannot aligns known gene sequences in existing databases (which are called “query sequences”) to assembled contigs (called “target sequences”) with minimap2. As some genes, such as *KIR2DL4* and *C4*, may have copy number variations, Immuannot allows one query sequence to be aligned to multiple locations on the target sequences. For each alignment, Immuannot calculates a gap-compressed identity as 𝑚/(𝑚 + 𝑑), where 𝑚 is the number of matching bases in the alignment and 𝑑 is the number of mismatching bases and gaps, regardless of gap length. Immuannot ranks alignments by the gap-compressed identity and then by the alignment length. It creates clusters of alignment by iterating alignments from the best to the worst. For a given alignment, Immuannot adds it to existing clusters if it overlaps with existing ones, or Immuannot creates a new cluster. Alignments that overlap with more than one existing clusters will be skipped because we only use full-length gene sequences and we also require high mapping rate (>90%). In the end, each cluster ideally represents a candidate gene. This greedy procedure assumes genes are well separated on the genome, which has not been an issue for HLA and KIR genes.

### Typing genes in the IPD-IMGT/HLA and IPD-KIR databases

At present, Immuannot uses IPD-IMGT/HLA database v3.52.0 and IPD-KIR v2.12 for typing. The two databases included 20,393 gene sequences and 38,684 partial/full CDS sequences. To fully leverage the IPD resources, only full length gene sequences are used for gene structure identification and all available CDSs are used for allele calling. Within each alignment cluster, Immuannot annotates gene structures and type alleles in the following three steps:

1. <u>Annotating gene structures</u>. Immuannot filters a gene sequence if less than 90% of the sequence is mapped or its gap-compressed identity is <97%. The remaining gene sequences are classified into four groups in decreasing priority: (a) with perfect alignment to the target contig; (b) with all CDS aligned and without gaps in CDS; (c) with all CDS aligned but with frameshifts; (d) with some CDS unaligned. If there are multiple sequences in one priority group, the gene sequence with fewest gap-compressed differences and then with longest alignment length is chosen as the template allele. The gene structure of template allele is lifted to the target contig to define the gene structure on the contig.
2. <u>Allele typing</u>. If the template allele in step (1) perfectly matches the target contig in full length, the template allele is concluded as the final allele call. Otherwise, CDS on target contig is extracted based on the annotated gene structures in step (1). The known CDS alleles in the IPD databases are aligned to the target CDS and are ranked by edit distance and then by alignment length. The partial CDS alleles are used only when the alignments cover the full length of query CDS and have no mismatches. Immuannot may report multiple closest alleles for a gene if their alignments to the contig have identical edit distance and alignment length.
3. <u>Naming novel alleles</u>. For a novel allele that is not present in the IPD databases, Immuannot may name it with a “new” keyword. For an HLA gene, if the best-matching alleles identified in step 2 come from multiple allele groups (i.e. the first fields in their names are different), Immuannot will call the allele as “HLA-X*new”, where “HLA-X” is the gene name. Otherwise, Immuannot finds the longest prefix among names of the best-matching alleles and assigns “new” to the second/third/fourth field, if the protein/CDS/non-coding sequence is affected, respectively.

### Typing C4 genes

Typing *C4* gene is easier than typing other HLA and KIR genes because Immuannot only attempts to distinguish between *C4A* and *C4B* and between the long form and the short form. It extracts the *C4A* transcript sequence on NG_011638.1 and aligns the sequence to assembled contigs in the splice mode to annotate *C4* gene structures. Immuannot distinguishes *C4A* and *C4B* based on four amino acid differences between the two genes on exon 26 (Blanchong et al. 2001; Sekar et al. 2016; Wang and Liu 2021). It considers the gene to have a long form if the length of intron 9 is longer than 5kb; otherwise Immuannot denotes the gene to have a short form.

### Population CDS divergence calculation

Gene divergence and diversity are calculated based on the CDS of each gene per sample. CDS sequences are extracted by gffread (Pertea and Pertea 2020), with the assemblies and our annotation gtf file as input. Then for each gene, CDSs are aligned by kalign3(Lassmann 2019) for each data panel (HPRC and CPC). Based on the aligned CDS, pairwise difference per sample is calculated as the average Hamming distance to the remained samples. The per gene diversity is calculated as the average of all pairwise differences of that gene.

### Construct HLA-DRB polygenetic tree

Full-length CDS of DRB1,3,4,5 are collected from HPRC, CPC, CHM13.0, GRCh38.p14, CN-T2T, IPD-IMGT/HLA, and IPD-IMGT/MHC, including 198 *HLA-DRB1*, 7 *HLA-DRB3*, 4 *HLA-DRB4*, 6 *HLA-DRB5* unique CDSs in total. DRB CDS from non-human primate species, chimpanzee (patr), crab-eating macaque (mafa), and rhesus monkey (mamu), are used as outgroup for tree construction, the most recent common ancestry of all outgroup sequences is set as the root in the phylogenetic tree. Kalign3 is used for the multiple sequence’s alignment and IQTree2 (Kalyaanamoorthy et al. 2017) is used for the tree construction with the option ‘-m MFP’ to select the best-fit model. Tree plot is generated with jstreeview.

### Estimate HLA gene deletion frequency in 1000 genome project data

To detect the deletion of HLA genes in 1000 genome data populations, we calculated the median read depth of the target gene region and 2000 randomly selected regions within chr6:2timb-5timb as the control that were supposed to be diploid. The read depth of a particular gene is normalized in two steps to reduce within- and between-sample variance. In step 1, the read depth of the gene is divided by half of the median read depth in the control regions of that individual. For a diploid individual without copy-number changes, this procedure would normalize the original read depth to a number close to two. In step 2, the normalized copy number from step 1 is divided by the median normalized copy number across all samples, assuming that the majority of the samples carry two copies of the targeted gene. This method turns out to work well to distinguish the copy number variation for the four-pseudo-gene deletion (*HTKU-del*) and *MICA* deletion (Supplemental Figure S9).

### Copy number estimation of T1K’s call

The copy number of a gene is inferred from the abundance estimate outputted by T1K. To infer the copy number, we first apply square root to abundance such as the resulting values approximately follow a normal distribution. We then use the smallest 30% of the transformed allele abundances to infer the mean and standard variance, denoted as μ and σ, respectively. This step can also incorporate the user-provided genes that are expected to have two copies. In this work, we assumed three HLA genes (*A*, *B*, and *C*) for HLA allele calling and four KIR framework genes (*3DL3*, *2DL4*, *3DP1*, and *3DL2*) for KIR allele calling. After obtaining the parameters of the normal distribution, our script will calculate the likelihood for each alleles copy number 𝑐 based on the formula:

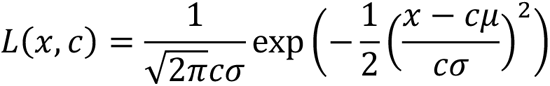

where x is the transformed abundance of an allele, c is the copy number, and this formular is based on the additive property of normal distribution. Then the copy number of this allele will be the value c that maximize the likelihood, e.g. 𝑎𝑟𝑔𝑚𝑎𝑥*_c_* 𝐿(𝑥, 𝑐).

## Data and Resources

HPRC assemblies: https://github.com/human-pangenomics/HPP_Year1_Assemblies

CPC assemblies: data access number PRJCA011422 at the National Genomics Data Center (https://ngdc.cncb.ac.cn).

GRC KIR contigs: source : Gentiank; access date: 2023-02-28; contig names: NT_187672.1, NT_187673.1, NT_187674.1, NT_187675.1, NT_187676.1, NT_187677.1, NT_187683.1, NT_187684.1, NT_187685.1, NT_187686.1, NT_187687.1, NT_187693.1, NW_016107300.1, NW_016107301.1, NW_016107302.1, NW_016107314.1, NW_016107313.1, NW_016107312.1, NW_016107311.1, NW_016107310.1, NW_016107309.1, NW_016107308.1, NW_016107307.1, NW_016107306.1, NW_016107305.1, NW_016107304.1, NW_016107303.1

Six HLA reference contigs: NCBI BioProject PRJNA764575

Immuannot: https://github.com/YingZhou001/Immuannot

Annotation data in this work: https://zenodo.org/records/8372992

## Acknowledgments

This project is supported by NHGRI grant R01HG010040 and U01HG010961 to H.L., and NIGMS grant P20GM130454 to Dartmouth (L.S).

## Author contributions

H.L. conceived and designed the project. Y.Z. implemented the software Immuannot. L.S. provided the script for copy number estimation of T1K’s call. Y.Z. conducted the analyses with input from L.S. and H.L. Y.Z., L.S. and H.L. wrote the manuscript. All the authors agree on the manuscript.

